# Representational dynamics of memories for real-life events

**DOI:** 10.1101/2022.02.18.480992

**Authors:** Olivier Jeunehomme, Rebekka Heinen, David Stawarczyk, Nikolai Axmacher, Arnaud D’Argembeau

## Abstract

The continuous flow of experience that characterizes real-life events is not recorded as such in episodic memory but is condensed as a succession of event segments separated by temporal discontinuities. To unravel the neural basis of this representational structure, we recorded real-life events using wearable camera technology and used fMRI to investigate brain activity during their temporal unfolding in memory. We found that, compared to the representation of static scenes in memory, dynamically unfolding memory representations were associated with greater activation of the posterior medial episodic network. Strikingly, by analyzing the autocorrelation of brain activity patterns at successive time points throughout the retrieval period, we found that this network showed higher temporal dynamics when recalling events that included a higher density of event segments. These results reveal the key role of the posterior medial network in representing the dynamic unfolding of the event segments that constitute real-world memories.

## INTRODUCTION

Decades of cognitive, neuropsychological, and neuroimaging research on episodic memory have deepened our understanding of the neurocognitive processes supporting the ability to remember past events.^1–5^ However, most studies have focused on memory for static stimuli (e.g., lists of words or pictures), so important questions remain about how the dynamic unfolding of events is represented. People make sense of the continuous stream of experience by segmenting it into meaningful units (i.e., events and sub-events), with this segmentation process playing a key role in the temporal structure of memory representations.^6–8^ Memories are composed of a succession of event segments (i.e., moments or slices of prior experience) that are organized in chronological order^9,10^ and usually include temporal discontinuities in the unfolding of events: Parts of the continuous flow of information that characterized the previous experience are omitted during event recall.^11–13^ This structure of memories allows one to mentally represent events in a temporally compressed form, such that it usually takes less time to remember an event than it took for this event to occur in the past.^14–18^ To date, however, the neural basis of this temporal structure of real-world memories remains unclear.

The neural basis of memories of real-life events has been mainly investigated in studies of autobiographical memory.^a^ Functional neuroimaging studies have shown that these memories are supported by a specific set of brain regions that includes the medial temporal lobes (e.g., hippocampus, parahippocampal gyrus), cortical midline structures (medial prefrontal cortex, precuneus, posterior cingulate cortex, and retrosplenial cortex), and lateral fronto-temporo-parietal areas.^20–24^ Within this core memory network, the hippocampus and medial parietal cortex have been associated with the process of “scene construction” (i.e., the generation, maintenance, and visualization of complex spatial scenes), which is considered one of the key components of past event recall.^25^ Supporting this view, it has been shown that episodic memory retrieval and the imagination of fictitious experiences are associated with overlapping patterns of brain activation, notably in the hippocampus, parahippocampal gyrus, and retrosplenial cortex.^26^ However, most studies on scene construction required participants to mentally represent static scenes (e.g., visualizing an old library),^27^ whereas, as noted above, memories for real-life events involve a dynamic succession of moments of prior experience.

A recent fMRI study has provided important insight into the neural mechanisms underlying the dynamic unfolding of naturalistic events in episodic memory.^28^ During the initial encoding of events from a movie, event segmentation processes were evidenced at multiple time scales throughout the cortical hierarchy: While early sensory regions represented relatively short-lived segments of experience, the posteromedial cortex and angular gyrus supported more extended episode representations that corresponded closely to event boundaries identified by human observers. Event boundaries in these higher-level areas triggered the hippocampus to encode the current situation model into episodic memory, which was later reinstated in the posteromedial cortex and angular gyrus during memory retrieval (see also refs.^29–31^). In a recent reanalysis of the same data set, Musz and Chen^32^ further showed that, relative to temporally precise utterances during recall, compressed event representations were associated with reduced neural reinstatement in the posteromedial cortex.

While these results present an important first step towards understanding the temporal dynamics of event perception and memory, the movies that were used differed from real-world events in important ways, notably because the events depicted were summarized using film editing techniques such as cuts, which affects memory representations.^33^ Thus, the dynamic unfolding of real-world memories remains to be elucidated. As noted above, these memories are not literal reproductions of past events, but instead consist of a succession of event segments that typically includes temporal discontinuities. The density of the segments of experience that constitute memories, and therefore the amount of temporal discontinuities within memories, can vary substantially across events, leading to more or less compressed representations.^17^ Here, we aimed to unravel the neural basis of this dynamic unfolding of the units of experience that constitute real-world memories.

We first sought to examine brain regions that are involved in remembering dynamic and temporally extended events as compared to static scenes. Memories were prompted using a paradigm that capitalizes on wearable camera technology to investigate the remembrance of real-life experiences.^12,34,35^ Specifically, participants performed a 40-min walk on a university campus during which they experienced a series of daily life events, while the content and timing of these experiences were recorded using a wearable camera^36^ (Figure 1A). Immediately after the walk, participants took part in an fMRI session in which they had to remember events that were cued by pictures that had been taken during their walk (Figure 1B). By contrasting these episodic memories to a control condition involving the mental representation of static scenes, we identified the brain regions that specifically underlie the recall of temporally unfolding events. We focused on the recall of real-world events rather than movies because the former involve a sense of agency that is absent from passive video viewing but may play an important role in structuring memory representations, given that a key function of episodic memory is to remember actions and goal-relevant information.^1,37,38^

**Figure 1:**
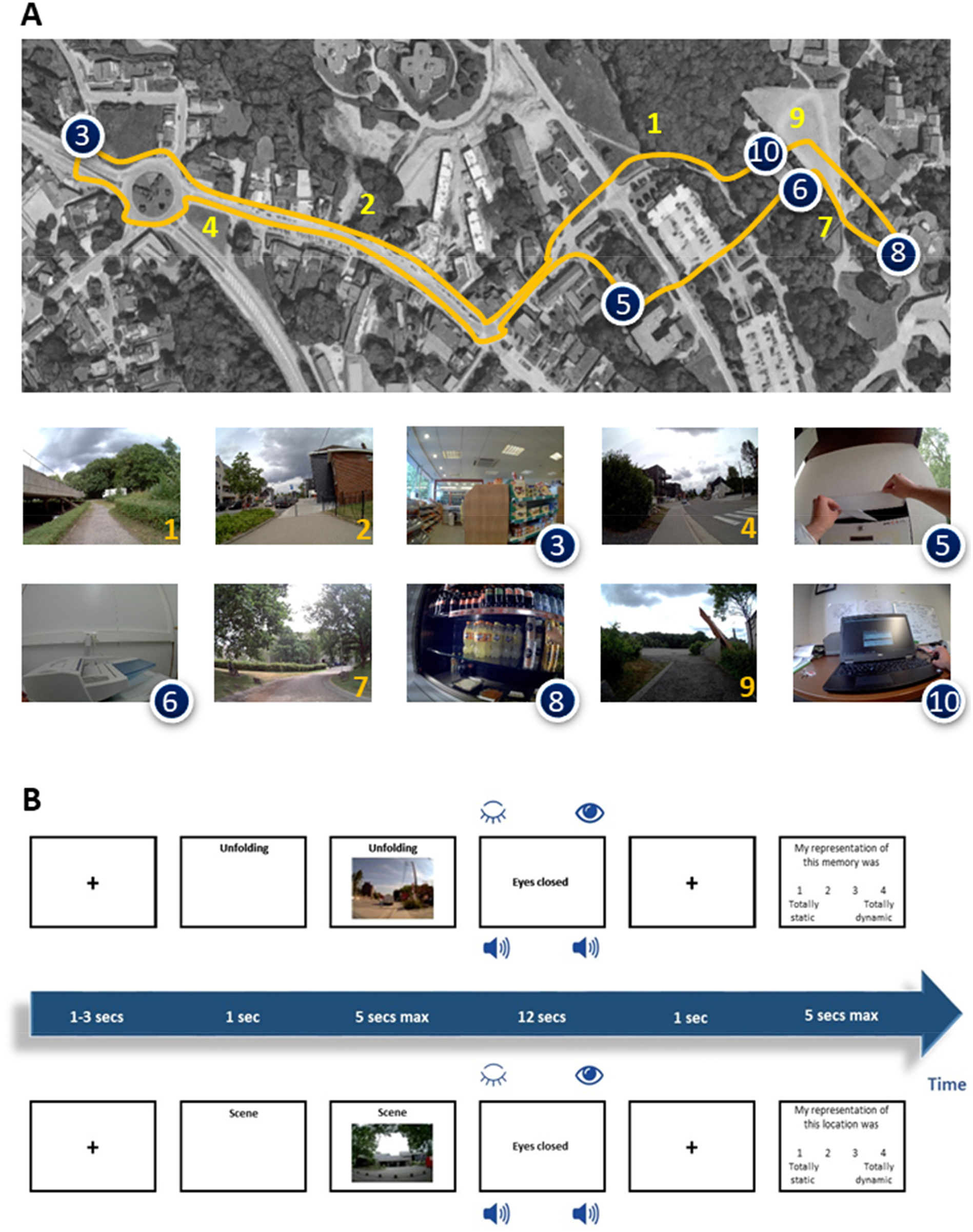
Experimental procedures. (A) Overview of the walk on the campus of the University of Liège. Locations in which participants performed an action are indicated by blue circles and the paths taken to go to these locations are indicated by yellow lines. For each event, an example of picture taken by the wearable camera that was used as cue for the episodic memory task is provided. (B) Trial structure in the episodic memory conditions (top panel) and in the control condition (bottom panel). Participants were first presented with a cue specifying the experimental condition. In the episodic memory conditions, a picture taken during the walk of the participant (corresponding to the beginning of an action or a spatial displacement) was then presented on the screen. As soon as they identified the moment of the walk depicted by the picture, participants were asked to remember the unfolding of the event, starting at the precise moment they identified on the picture. In the control condition (i.e., scene condition), participants were asked to imagine the visual scene depicted by the picture (which was not part of their walk). For each condition, participants were asked to close their eyes and to press a response key once they started to perform the task; they did the task with eyes closed until the presentation of a sound signal (which was presented 12 seconds after their button press). Finally, each trial ended with the presentation of a Likert scale on which participants rated the temporal dynamics of their mental representation (i.e., static vs. dynamic). In total, 20 trials were presented for each condition. FMRI analyses focused on the time interval between the key press and the end of the 12 second period, thus corresponding to the phase of mental replay.

The central aim of the current study was then to compare the neural correlates of episodic memories that are associated with different event compression rates. Previous studies have shown that the density of the units of experience that constitute real-world memories, and therefore the temporal compression of events, is not constant but varies with the content of remembered events. In particular, memories for actions contain a higher density of experience units than memories for spatial displacements.^12,13,17^ Indeed, spatial displacements involve a relatively simple structure that is primarily organized in terms of changes in direction (i.e., turns), whereas actions typically have a more complex and fine-grained structure in which event segments correspond to changes in goals and sub-goals.^39^ This difference in the density of event segments should be reflected in the dynamics of neural activity when remembering the unfolding of events.

Our main hypothesis is that the posterior medial network, which is thought to support the representation of event models,^40–42^ plays a key role in representing the temporal structure of memories. Indeed, regions such as the retrosplenial cortex, posterior cingulate cortex, precuneus, and angular gyrus might contribute to the integration of successive changes in experience when remembering the unfolding of events, and might also maintain and elaborate integrated event representations.^41^ To test these hypotheses, we not only examined brain activity over the entire duration of the retrieval period using classical univariate analyses, but also investigated to what extent multivariate patterns of brain activity change across time during the unfolding of events in memory. To the extent that memories for actions contain a higher density of experience units, we expected that they would be associated with higher temporal dynamics (i.e., lower autocorrelations) of brain activity patterns at successive time points during the retrieval period.

## RESULTS

### Behavioral results

Participants successfully identified the presented pictures for nearly all trials (*M* = 98.24%, 95% CI [98.11%, 98.36%]). As expected, the ratings obtained for each trial during scanning (see STAR Methods) indicated that the mental representations formed in the two episodic memory conditions were dynamic, whereas the mental representations formed in the control condition were static (see Table 1 for mean ratings). Ratings were not significantly different between the two memory conditions (i.e., action and spatial displacement), *t* = 1.74, *p* = .108, *ξ* = .34.

**Table 1.**
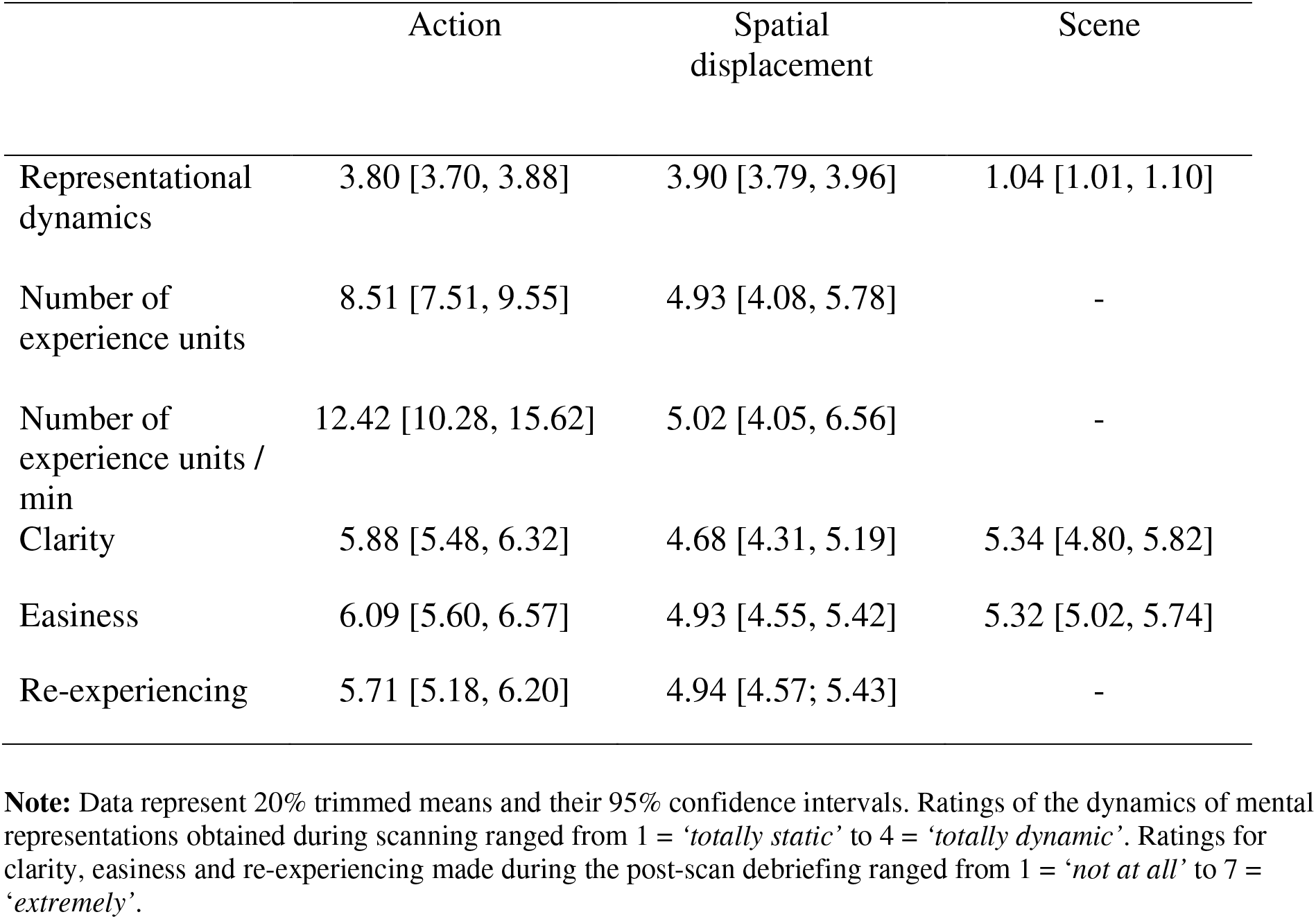
Behavioral data for the episodic memory (action, spatial displacement) and control (scene) conditions

|  | Action | Spatial displacement | Scene |
| --- | --- | --- | --- |
| Representational dynamics | 3.80 [3.70, 3.88] | 3.90 [3.79, 3.96] | 1.04 [1.01, 1.10] |
| Number of experience units | 8.51 [7.51, 9.55] | 4.93 [4.08, 5.78] | - |
| Number of experience units / min | 12.42 [10.28, 15.62] | 5.02 [4.05, 6.56] | - |
| Clarity | 5.88 [5.48, 6.32] | 4.68 [4.31, 5.19] | 5.34 [4.80, 5.82] |
| Easiness | 6.09 [5.60, 6.57] | 4.93 [4.55, 5.42] | 5.32 [5.02, 5.74] |
| Re-experiencing | 5.71 [5.18, 6.20] | 4.94 [4.57; 5.43] | - |

**Note:** Data represent 20% trimmed means and their 95% confidence intervals. Ratings of the dynamics of mental representations obtained during scanning ranged from 1 = ‘*totally static*’ to 4 = ‘*totally dynamic*’. Ratings for clarity, easiness and re-experiencing made during the post-scan debriefing ranged from 1 = ‘*not at all*’ to 7 = ‘*extremely*’.

The verbal descriptions of memories obtained immediately after the scanning session showed that participants re-experienced a succession of experience units when recalling the events (see STAR Methods). Memories of actions contained more experience units than memories of spatial displacements, *t* = 7.29, *p* < .001, *ξ* = .85 (Table 1). To assess the density of recalled experience units, we computed the number of reported units per minute of the actual event duration (see STAR Methods). In line with previous studies,^12,13,17^ the density of experience units was higher when recalling actions than spatial displacements, *t* = 6.77, *p* < .001, *ξ* = .88. As shown in Table 1, all memory representations were judged as clear, easy to recall/construct, and associated with a subjective feeling of re-experiencing the events.

### fMRI results

#### Univariate activity associated with episodic memories versus scene representations

Although our main interest was in the temporal dynamics of brain activity patterns during event recall, we first sought to check that our novel paradigm would engage the brain network that has been associated with memories for real-life events in previous studies. To do so, we conducted univariate analyses to identify brain regions within the autobiographical memory network (see STAR Methods) that showed higher activity when recalling the unfolding of events compared to static scene representations (see Figure 2A, Table 2). Results showed that the recall of actions (action > scene) was associated with increased activation in bilateral parietal areas, including the precuneus, posterior cingulate, retrosplenial cortex, and angular gyrus. Moreover, we found higher activity in the superior/middle frontal gyrus, the posterior middle temporal gyrus, middle occipital cortex, and cerebellum. When looking at memories of spatial displacements (spatial displacement > scene), similar activations were observed in medial parietal areas (i.e., precuneus, posterior cingulate, and retrosplenial cortex). Additional clusters of activation were detected in the right angular gyrus/middle occipital cortex, and bilaterally in the middle frontal gyrus. Figure 2A also illustrates overlapping activations associated with each contrast, which show that the two types of episodic memories were associated with common activation in medial parietal areas, including the precuneus, posterior cingulate cortex, and retrosplenial cortex.

**Figure 2.**
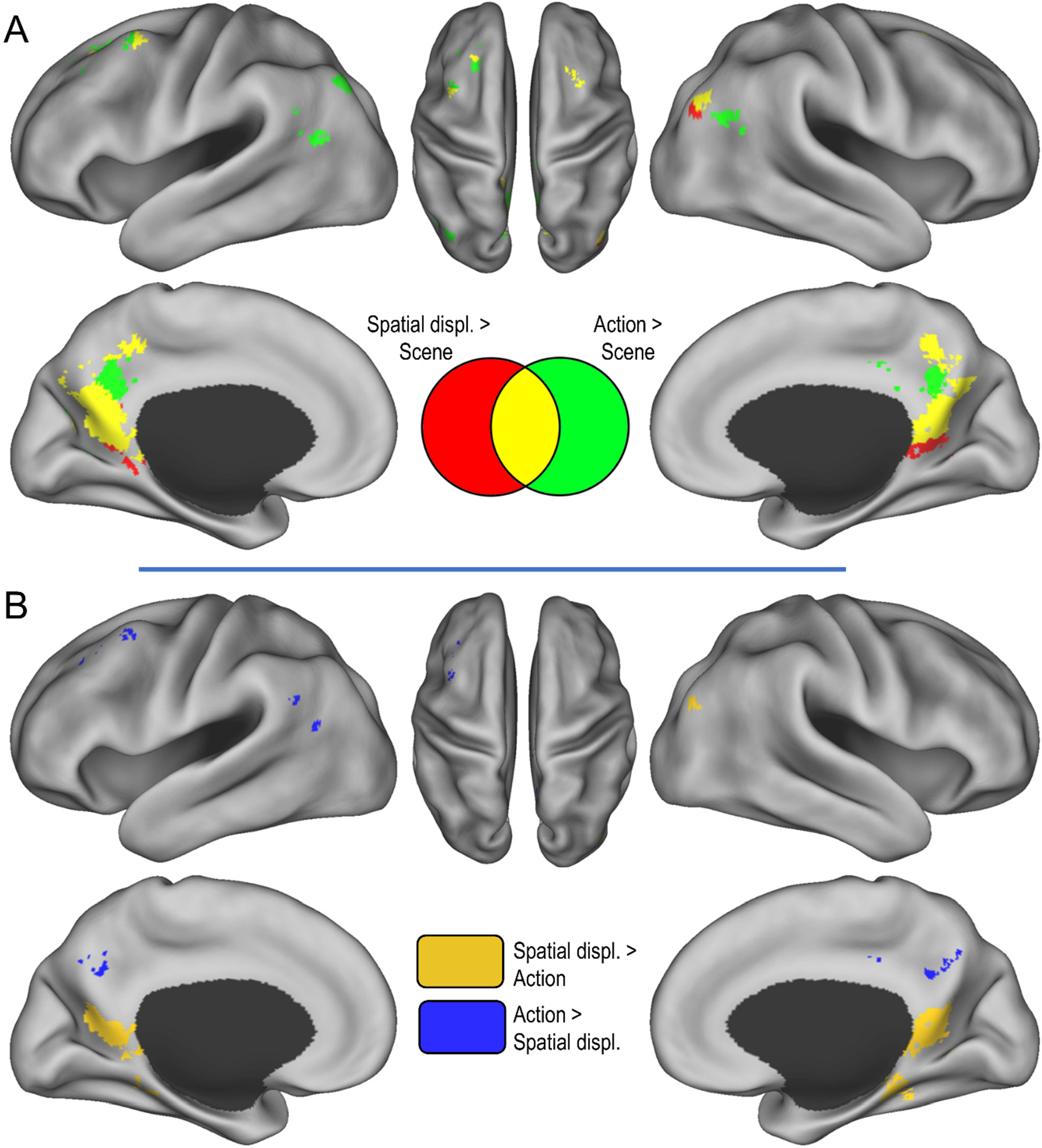
Brain regions activated when recalling the unfolding of events (univariate analyses). (A) Regions that were more activated when recalling actions and spatial displacements compared to the scene construction condition. (B) Regions showing differential activity when recalling actions and spatial displacements. Activations are displayed at *p* < .001 (uncorrected) within the autobiographical network mask.

**Table 2.** Brain regions showing increased activity for episodic memories as compared to scene representations

| Brain regions | Side | MNI coordinates |  |  | Cluster Size<br>(n° voxels) | t |
| --- | --- | --- | --- | --- | --- | --- |
|  |  | X | Y | Z |  |  |
| ***Actions > scenes*** |  |  |  |  |  |  |
| Precuneus/PCC/RSC | R | 12 | -62 | 36 | 2111 | 11.46 |
|  | R | 8 | -54 | 44 |  | 9.93 |
|  | R | 8 | -38 | 26 |  | 8.35 |
|  | L | -12 | -60 | 20 |  | 8.79 |
|  | R | 12 | -62 | 28 |  | 10.99 |
| Middle cingulate gyrus | L | -2 | -32 | 40 | 28 | 6.71 |
| Middle/superior frontal gyrus | L | -40 | 10 | 54 | 84 | 6.40 |
|  | R | 30 | 14 | 58 | 35 | 5.85 |
| Middle frontal gyrus | L | -24 | 30 | 48 | 172 | 4.73 |
| AG | R | 44 | -68 | 36 | 165 | 5.66 |
|  | L | -58 | -56 | 26 | 21 | 4.40 |
| MTG/AG | L | -48 | -54 | 20 | 66 | 5.64 |
| Middle OC | L | -40 | -72 | 38 | 64 | 5.38 |
| MTG | R | 62 | -52 | 10 | 20 | 5.18 |
| Cerebellum | L | -6 | -54 | -40 | 28 | 4.18 |
| ***Spatial displacements > Scenes*** |  |  |  |  |  |  |
| Precuneus/PCC/RSC | L | -10 | -58 | 18 | 1983 | 11.03 |
|  | R | 12 | -52 | 20 |  | 8.64 |
|  | R | 4 | -44 | 12 |  | 7.75 |
|  | R | 8 | -64 | 44 |  | 7.48 |
| AG/Middle OC | R | 42 | -70 | 34 | 92 | 5.37 |
| Middle frontal gyrus | R | 28 | 16 | 54 | 31 | 4.91 |
|  | L | -34 | 6 | 60 | 40 | 4.72 |
|  | L | -38 | 8 | 54 |  | 4.40 |
|  | L | -22 | 26 | 42 | 23 | 4.17 |

**Note:** Coordinates refer to brain regions identified in the SPMs thresholded at $p < .001$ (uncorrected) that were statistically significant (at $p < .05$ ) using SVC (see STAR Methods). PCC = Posterior cingulate cortex; RSC = Retrosplenial cortex; OC = Occipital cortex; AG = Angular gyrus; MTG = Middle temporal gyrus.

Next, we compared the two episodic memory conditions (actions vs. spatial displacements; see Figure 2B, Table 3). The retrieval of actions was associated with higher activity in the precuneus, left middle frontal gyrus, right middle temporal gyrus, middle cingulate cortex, and angular gyrus compared to the retrieval of spatial displacements. The reverse contrast (spatial displacements > actions) revealed increased activations in the retrosplenial cortex, the right parahippocampal gyrus, and the right middle occipital gyrus.

**Table 3.** Brain regions showing differential activity for actions vs spatial displacements

| Brain regions | Side | MNI coordinates |  |  | Cluster Size<br>(n° voxels) | t |
| --- | --- | --- | --- | --- | --- | --- |
|  |  | X | Y | Z |  |  |
| **Action > Spatial displacement** |  |  |  |  |  |  |
| Middle frontal gyrus | L | -32 | 26 | 36 | 34 | 6.25 |
| Precuneus | L | -2 | -62 | 38 | 131 | 5.51 |
| MTG | R | 64 | -48 | 6 | 22 | 5.45 |
| Middle cingulate gyrus | R | 2 | -26 | 38 | 21 | 5.39 |
| AG | L | -58 | -54 | 30 | 24 | 4.82 |
| **Spatial displacement > Action** |  |  |  |  |  |  |
| RSC | L | -12 | -52 | 12 | 483 | 9.44 |
|  | R | 12 | -46 | 6 | 363 | 7.84 |
| PHG/FG | R | 20 | -34 | -10 | 170 | 7.68 |
| Middle occipital gyrus | R | 42 | -74 | 26 | 21 | 4.28 |

**Note:** Coordinates refer to brain regions identified in the SPMs thresholded at $p < .001$ (uncorrected) that were statistically significant (at $p < .05$ ) using SVC (see STAR Methods). RSC = Retrosplenial cortex; MTG = Middle temporal gyrus; PHG = Parahippocampal gyrus; FG = Fusiform gyrus; AG = Angular gyrus.

#### Patterns of brain activity across recall conditions

We next examined whether distributed patterns of brain activity in the autobiographical memory network (using the same mask as for the univariate analyses) reflected the content of memories – i.e., whether these patterns differed systematically between experimental conditions. Using representational similarity analysis, we compared all recall trials of the three experimental conditions (actions, spatial displacements, scenes) with trials of the same condition as compared to trials of other conditions. A repeated measures ANOVA revealed a significant main effect of within versus between similarities (*F*(1,20) = 34.58, *p* = <0.0001), which showed significantly higher within condition correlations compared to between condition correlations for all three conditions (Figure 3A). There was no main effect of condition (action/spatial displacement/control; *F*(2,40) = 1.21, *p* = 0.308) and no interaction (*F*(2,40) = 1.49, *p* = 0.238).

**Figure 3.**
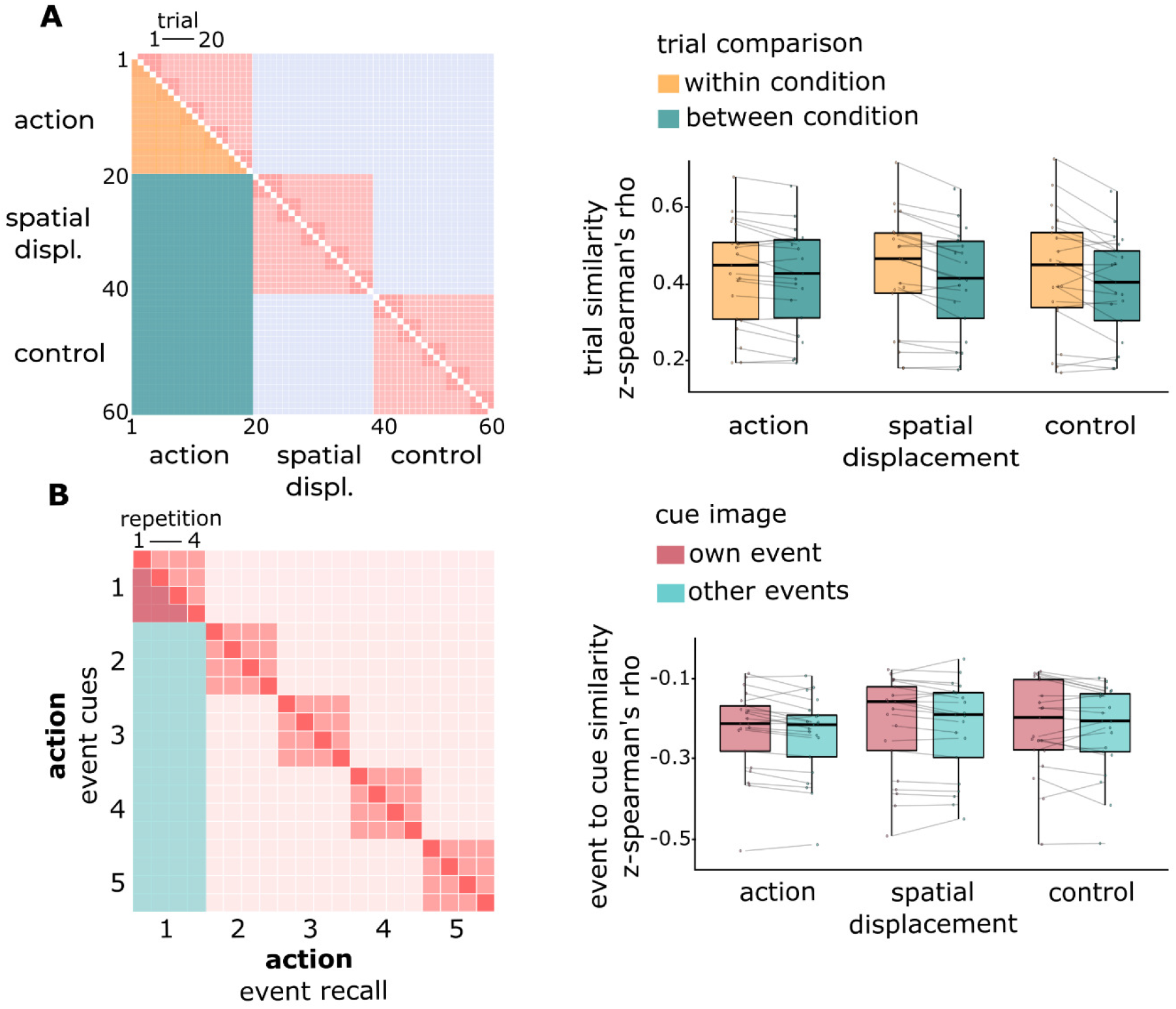
Content-specific memory representations in the autobiographical memory network. (A) For each condition (here illustrated for actions on the left panel), we compared trials within each condition (yellow) versus all other trials (green). Distributed patterns of neural activity were more similar when recalling events from the same condition than events from different conditions. This effect is observed across all experimental conditions (right panel). (B) For each event in each condition (here illustrated for actions, event 1, on the left panel), we compared the similarity between the recall period and the image cue of the same event (red) to image cues of other events (blue), excluding same trial similarities (diagonal). Recall patterns were more similar to their event-specific image cue than to image cues of other events in the same condition (right panel). This effect is observed across all experimental conditions. Dots show descriptive means for each subject; individual observations are linked with lines.

One may argue that these higher similarities for trials within conditions might be caused by similarities in types of neural processes engaged during recall (e.g., different amounts of memory search processes) rather than recalled contents. To address this issue, we further computed the similarity of each of the four recall repetitions of an event with the activity pattern during their image cue and compared it to the similarity with the image cues of the other events of the same condition (Figure 3B). Since similarities within trials tend to be higher than between trials, we calculated, for each event, neural activity during the recall phase in trial m with activity during the image cues in trials n≠m of the same event and compared this to correlations with image cues of the other events in the same condition. A repeated measures ANOVA showed a significant main effect of cue type (own vs. other events; *F*(1,20) = 17.05, *p* = 0.0005), no main effect of condition (*F*(2,40) = 1.51, *p* = 0.235), and no interaction (*F*(2,40) = 0.67, *p* = 0.499).

Overall, these initial analyses suggest that voxel patterns reflected remembered contents and can thus be used to analyze the temporal dynamics of episodic memories, which we did in the next step.

#### Temporal dynamics of brain activity patterns during event recall

We computed temporal autocorrelations between successive patterns of brain activity as a measure of the temporal dynamics of memories (see Figure 4A). A searchlight analysis revealed higher temporal dynamics (i.e., lower autocorrelations) during the recall of episodic memories compared to the representation of static scenes. Compared to the control condition, the recall of actions was associated with higher temporal dynamics in the precuneus, posterior cingulate cortex, angular gyrus, and lingual gyrus (*t*(20)= 10.8; *p*_corr_ < 0.05, FWE corrected using TFCE; Figure 4B/C). This extended cluster also included the middle frontal gyrus, frontal pole, middle temporal gyrus, and precentral gyrus. Compared to the control condition, the recall of spatial displacements was associated with higher temporal dynamics in the posterior cingulate cortex, cuneus, angular gyrus, and lateral occipital cortex (*t*(20)= 5.48; *p*_corr_ < 0.05, FWE corrected using TFCE; Figure 4B/C).

**Figure 4.**
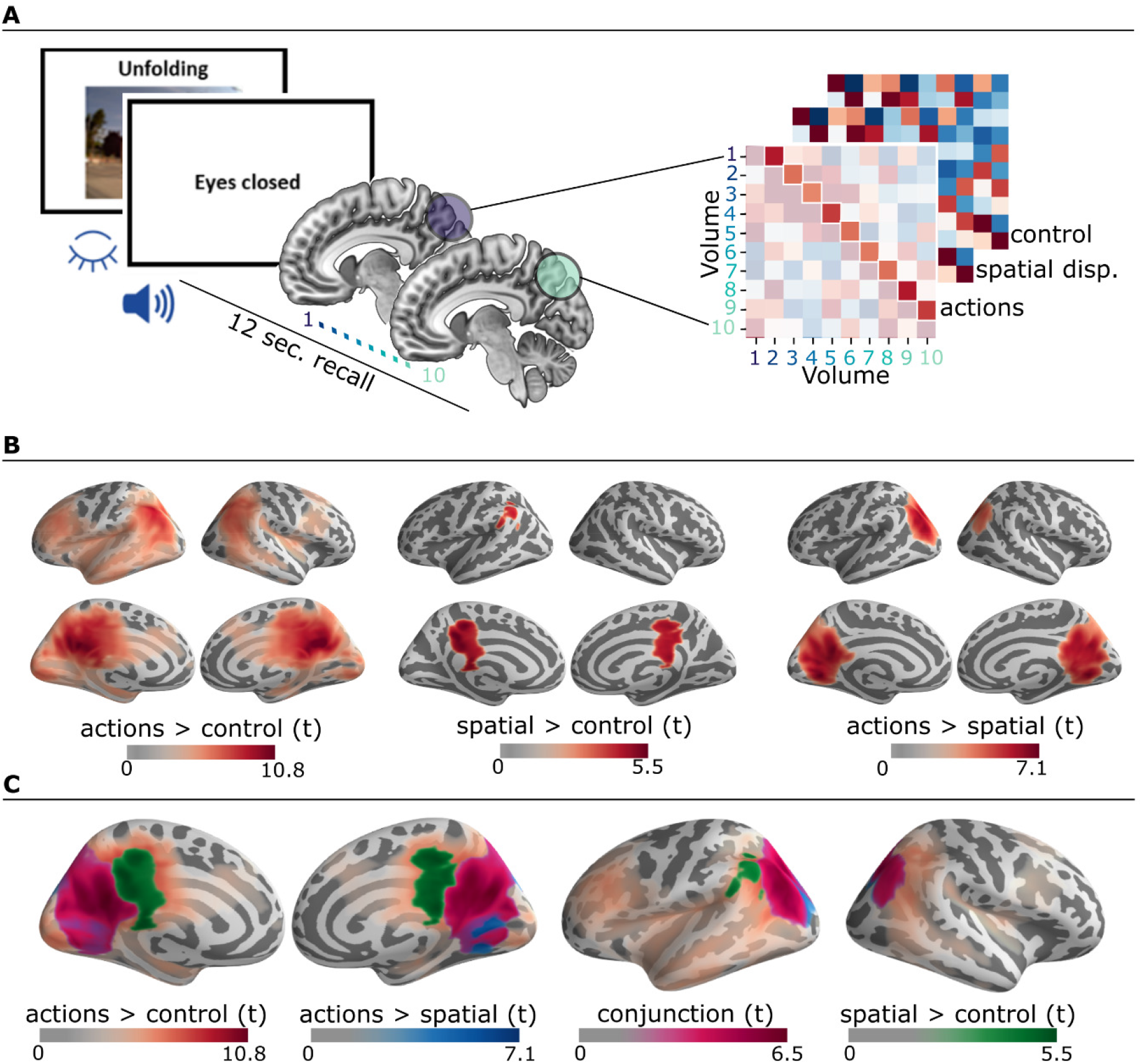
Temporal dynamics of patterns of brain activity during the recall period. (A) Within each searchlight sphere (first volume blue circle), we correlated 10 consecutive volumes (starting with the fifth volume after the recall onset to account for the delayed hemodynamic response function), resulting in a 10×10 spearman rho correlation matrix. From this matrix we extracted the mean of the first off-diagonal (i.e., correlations of patterns between all pairs of temporally adjacent volumes). This procedure was applied to all trials of all conditions and for all sphere locations, resulting in three brain maps (one for each condition) indicating the averaged autocorrelation as an (inverse) measure of the temporal dynamics of episodic recall. (B) The searchlight analysis revealed significantly higher temporal dynamics for the action condition compared to the control and spatial displacements conditions. Spatial displacements showed higher temporal dynamics compared to the control condition. (C) The conjunction of higher temporal dynamics for actions compared to both the spatial displacements and control conditions is shown in purple (The conjunction of spatial displacements vs. control and of action vs. control is not shown as it overlaps completely with the action vs. control contrast.).

Next, we compared the temporal dynamics of the two types of episodic memories and found higher temporal dynamics for the recall of actions than spatial displacements in the lateral occipital cortex, precuneus, posterior cingulate, cuneus, and angular gyrus (*t*(20) = 7.06; *p*_corr_ < 0.05, FWE corrected using TFCE, Figure 4B/C). None of the opposite contrasts showed any significant differences.

In addition to this whole-brain analysis, we conducted a second searchlight analysis within the autobiographical network mask that we used for the univariate analyses (Figure 5). This analysis revealed significantly higher temporal dynamics during the recall of actions compared to the control task within the precuneus and posterior cingulate (cluster 1: *t*(20) = 12.3; *p*_corr_ < 0.05, FWE corrected using TFCE), middle temporal gyrus (cluster 2: *t*(20) = 5.52; *p*_corr_ < 0.05, FWE corrected using TFCE), superior frontal gyrus (cluster 3: *t*(20) = 4.48; *p*_corr_ < 0.05, FWE corrected using TFCE), lateral occipital cortex and angular gyrus (cluster 4: *t*(20) = 4.15; *p*_corr_ < 0.05, FWE corrected using TFCE). We also found higher temporal dynamics for the action condition compared to the recall of spatial displacements within one cluster including the precuneus, lingual gyrus, cuneus, angular gyrus, and the lateral occipital cortex (*t*(20)= 8.75; *p*_corr_ < 0.05, FWE corrected using TFCE). The contrast between the spatial displacement and control conditions did not yield significant differences within the autobiographical network mask.

**Figure 5.**
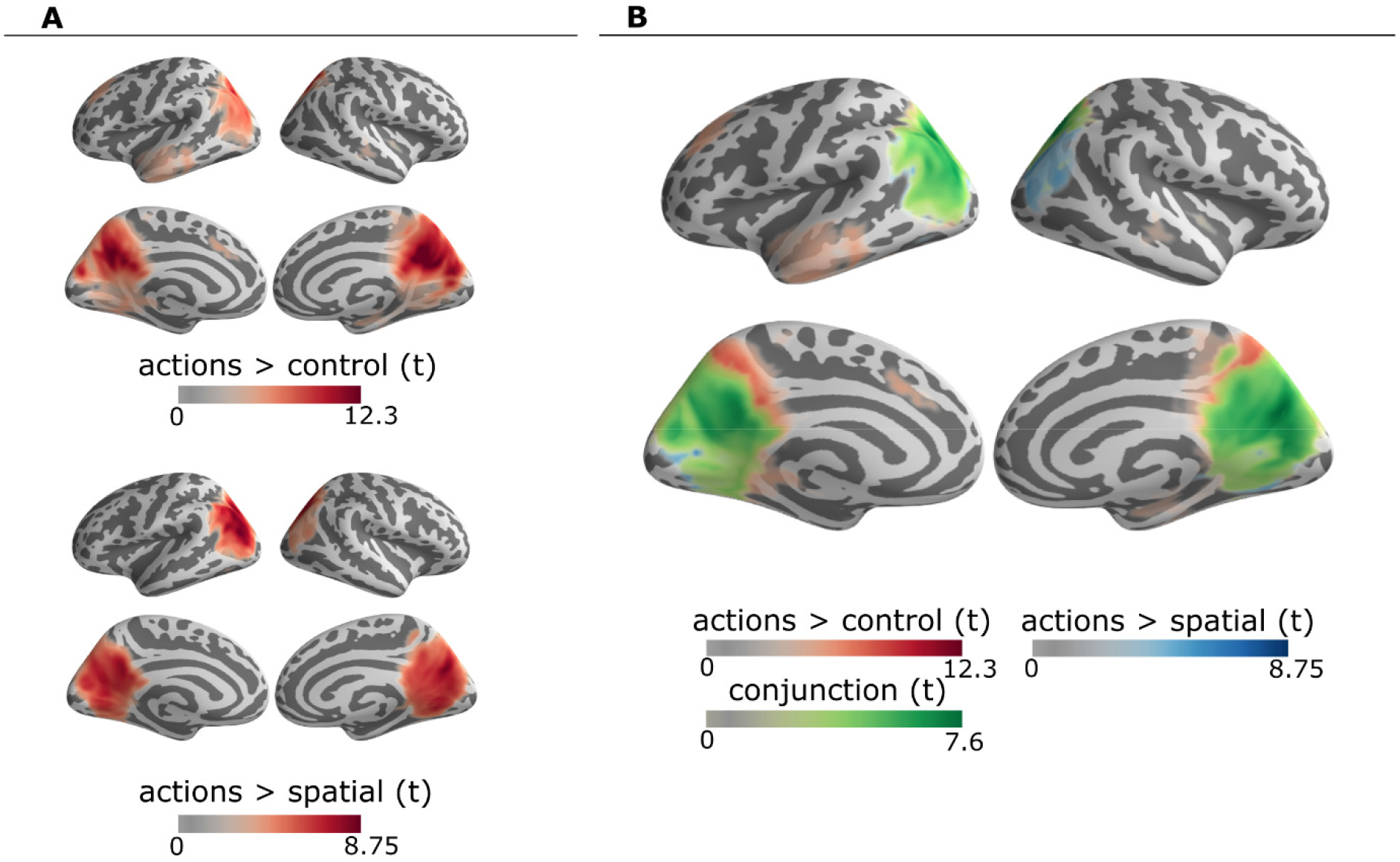
Searchlight analysis of temporal dynamics within the autobiographical network mask. (A). This analysis revealed significantly higher temporal dynamics for the action condition compared to both the control and the spatial displacement condition. (B) The conjunction of higher temporal dynamics for actions compared to spatial displacements and compared to the control condition is shown in green.

## DISCUSSION

The aim of our study was to better understand the neural mechanisms underlying the dynamic unfolding of events within episodic memory representations. Compared to mental representations of static scenes, memories of dynamic events elicited greater activation in medial parietal areas (including the precuneus, posterior cingulate cortex, and retrosplenial cortex) and in the angular gyrus. Furthermore, when investigating the autocorrelation of patterns of brain activity throughout the retrieval period, we found that these posterior brain regions showed greater temporal dynamics when remembering dynamic events compared to static scenes. These effects were more pronounced for memories that included a higher density of experience units (i.e., memories for actions compared to spatial displacements). Interestingly, the posterior brain regions that were identified as showing higher temporal dynamics without a priori location constraints (i.e., in the whole-brain searchlight analysis) overlapped with regions previously identified as part of the autobiographical memory network.

Memories for real-life events depend on an extended fronto-temporo-parietal network, but the exact functions of specific regions within this network are not fully understood.^20–24^ Medial parietal areas, including the precuneus, posterior cingulate, and retrosplenial cortex, are central areas of this network^24^ that share reciprocal connections^43^ and have been referred to as the posterior medial episodic system.^28,40,41^ A first contribution of the present study is to show that this posterior network is more active when recalling the dynamic unfolding of events compared to mentally representing static scenes. Remembering the unfolding of events involves the retrieval of moments or slices of prior experience (each moment itself consisting of a bounded set of various event features) that need to be sequenced and integrated into a stable and coherent representation.^1,44^ The posterior medial episodic network is considered to play a central role in the elaboration and use of event models (i.e., models that specify the spatial, temporal, and causal relationships between entities within a particular context),^40–42^ and may thus contribute to organize and integrate the sequence of experience units that represent the unfolding of events during episodic recall.

Another important aim of this study was to compare the neural correlates of episodic memories that are composed of different densities of experience units. Consistent with previous findings,^12,13,17^ our behavioral results showed that the density of experience units was higher when remembering actions compared to spatial displacements. Actions typically have a more complex and fine-grained segmental structure than spatial displacements,^39^ which is manifest when mentally representing the unfolding of prior experience during episodic recall. At the neural level, we found that some regions of the posterior medial episodic network, including the precuneus, were more active when remembering actions compared to spatial displacements. Furthermore, and most importantly, we observed higher temporal dynamics within patterns of activity in the precuneus, as well as the posterior cingulate and occipital cortex, when remembering the course of action events than when remembering spatial displacements or mentally representing static scenes. The precuneus has been associated with central aspects of autobiographical memory retrieval, including visuo-spatial imagery, first-person perspective taking, and an experience of agency.^43^ Interestingly, recent findings suggest that the precuneus plays a causal role in the dynamics of memory retrieval,^45^ and may notably contribute to the temporal structure and order of moments of past events.^46,47^ The higher temporal dynamics of activity patterns in the precuneus and other posterior regions that we observed suggest that these regions are sensitive to changes in recalled content during the retrieval period and may thus contribute to implementing and organizing transitions between experience units when representing the temporal unfolding of past events. This interpretation fits with previous research showing that patterns of activity in posterior medial regions are less similar across than within segments of a video episode.^28^

We also found that the angular gyrus was more active and showed greater temporal dynamics when remembering actions compared to spatial displacements. Several studies have established relationships between activity in the angular gyrus, especially left-lateralized, and the amount of recollected information and strength of recollection.^48–51^ Although the exact role of this region has been a matter of considerable debate,^52,53^ a recent unifying framework has proposed that the angular gyrus supports the integration and representation of the multimodal contextual details that constitute a mnemonic representation and promote vivid re-experiencing.^54^ The present findings suggest that this integration process contributes to representing the dynamic unfolding of experience units within episodic memory representations.

Compared to the recall of actions, the recall of spatial displacements was associated with increased activity in the retrosplenial cortex and parahippocampal gyrus. These regions are well known for their role in spatial navigation, and notably respond to relevant spatial elements of the environment such as landmarks.^55^ Interestingly, the parahippocampal gyrus coordinates observed in the present study overlap with those reported in a previous study on the role of the parahippocampal place area on scene perception.^56^ Evidence suggests that the parahippocampal place area and the retrosplenial cortex might play complementary functions in building a coherent and detailed representation of a scene across several viewpoints.^56,57^ Specifically, it has been proposed that when viewpoint changes occur continuously in a scene, the parahippocampal place area is sensitive to changes in physical details across each view, thereby representing scenes as a succession of individual representations of each view rather than a unique and integrated representation of multiple views. On the other hand, the retrosplenial cortex links the current and previous views of the same scene, creating an integrated representation of this scene over changes in visual perspective.^56^ In the current study, it is therefore possible that the parahippocampal gyrus and the retrosplenial cortex were more active when remembering the unfolding of spatial displacements compared to actions because the former implied more important updating and integration of successive views of the surrounding environment when mentally travelling from one particular moment of past experience to the next.

The critical role of the hippocampus in the retrieval of episodic memories is well established.^2^ This region is considered to play an important role in binding the various elements that constitute memories, including information about the sequence of events.^58–60^ Therefore, one could have expected that, in the present study, the hippocampus would be differentially activated when remembering events with different densities of experience units, but this is not what we found. We have no ready explanation for this null result, but it is worth noting that in previous work on memory for dynamic events, hippocampal activity was observed during encoding but not during event recall.^28^ One possible explanation could be that hippocampal activity during retrieval is transient in nature^61^ and thus was not captured by our analysis. Alternatively, the episodic segments supported by the hippocampus may be substantially longer (~1 minute)^28^ than the relatively short recall periods that we analyzed (12 seconds). A third possibility is that the lack of hippocampal activity in the present study is related to the fact that participants were highly familiar with the university campus where the events took place. Indeed, it has been found that the hippocampus is involved in memory for a recently learned, but not a highly familiar environment.^62^ In addition, in the present study, the events were recalled four times, which may have further increased their familiarity. To investigate the possible effect of repetitions, we conducted an additional RSA comparing the similarities of each trial to all other repetitions of the same event. This analysis indicated that neural activity patterns remained overall stable across trial repetitions for the action, spatial displacement, and control conditions (see Supplemental Information). In any case, future studies should investigate the contribution of the hippocampus to the temporal dynamics of memories in more detail.

We investigated the temporal dynamics of brain activity as a function of the density of recalled experience units by comparing memories for two types of events that have been repeatedly shown to differ in terms of event compression rates (i.e., actions vs spatial displacements).^12,17^ However, it should be noted that, in addition to these differences in the density of recalled units, other event characteristics might also have contributed to the observed differences in brain activity. For example, the action events mainly involved interactions with focal objects in indoor environments, whereas spatial displacements involved larger-scale outdoor environments.

Finally, it should also be noted that we do not consider the compression rates of events in episodic memory to be fixed. On the contrary, compression rates can be voluntarily modulated according to retrieval goals.^58^ For example, when people scan their memory for a particular piece of information, they skip more event segments and thus compression rates are higher than when they try to mentally relive the events.^18,63^ An interesting avenue for future research would be to use the method we developed for analyzing the temporal dynamics of brain activity patterns to investigate the neural basis of this flexible modulation of event compression during episodic retrieval.

In sum, the present study provides first insights into the neural correlates of the dynamic succession of experience units that constitute memories for real-life events. Our results show that remembering the unfolding of events activates posterior brain structures (including the precuneus, posterior cingulate cortex, retrosplenial cortex, and angular gyrus) that support event models and may contribute to the representation and integration of episodic details to form coherent representations of the succession of moments of prior experience. Furthermore, an important finding of the current study is that changes in patterns of activity within these regions reflect differences in the density of recalled experience units, suggesting that the posterior medial network plays an important role in the temporal dynamics of event recall and may notably contribute to implement transitions between the units of experience that represent the unfolding of events.

### Limitations of the study

From a methodological perspective, it would be interesting in future studies to compare our method of analysis of the temporal dynamics of brain activity patterns with other methods that detect state boundaries in activity patterns using hidden Markov models (HMM)^28^ or greedy state boundary search (GSBS).^64^ HMM and GSBS were not applicable to our data for several reasons. First, these methods have been applied to fMRI recordings ranging from 8 minutes^64^ up to 45 minutes,^28^ whereas recall trials in our study were limited to 12 seconds. State boundaries might not be detectable within such short trials, as Geerligs et al.^64^ found that the algorithms start to struggle with short state durations. Second, HMM and GSBS methods are applied to group data, detecting state boundaries across participants. The real-life events experienced by our participants were not completely comparable across participants, contrary to the standardized movies used in the aforementioned studies. In future studies, applying state detection algorithms on the recall of real-life events while taking these issues into account would be a promising next step for further characterizing the representational dynamics of memories.

Another limitation of the present study is that, although the memories that we instigated involved events happening in the real world, these events were constrained to be standardized across participants. This was done to investigate events that clearly differed on the density of recalled experience units (i.e., actions vs. spatial displacements), based on previous behavioral findings.^12,13^ In future studies, it would be interesting to use a similar paradigm with more idiosyncratic events that participants would record in their own life circumstances and to investigate whether and how various event features (e.g., emotional significance, self-relevance, familiarity) affect the representational dynamics of memories.

## Acknowledgments

This work was supported by the University of Liège (Fonds spéciaux – Crédits sectoriels no. 9893). O.J., D.S., and A.D. are supported by the Fund for Scientific Research (F.R.S.-FNRS), Belgium.

## Author Contributions

Conceptualization: A.D. and O.J.; Methodology: A.D. and O.J.; Software: R.H.; Formal Analysis: R.H. and O.J., under the supervision of N.A. and A.D.; Investigation: O.J. and D.S.; Writing – Original Draft: A.D., O.J. and R.H.; Writing – Review & Editing: N.A. and D.S. Visualization: R.H. and D.S.; Supervision: A.D. and N.A.; Funding acquisition: A.D.

## METHODS

### Participants

Twenty-one right-handed students (13 males and 8 females; mean age = 22.29 years, SD = 2.12, range = 18-27 years) from the University of Liège took part in this study. All participants were highly familiar with the university campus where the pre-scan walk took place. None of them reported a current use of psychoactive medication or a history of a neurological or psychiatric disorder. Five additional participants were tested but were excluded from the analyses due to noncompliance with the task, either during the pre-scan walk (one participant) or in the scanner (four participants). A formal power analysis was not performed because we did not find any prior study that involved a similar analysis as our main analysis of interest (i.e., examining autocorrelations between patterns of brain activity across time). Thus, our sample size was selected to be comparable to previous studies investigating memory for dynamic stimuli.^28^ All participants gave their written informed consent to take part in the study, which was approved by the ethics committee of the Medical School of the University of Liège and was performed in accordance with the ethical standards described in the Declaration of Helsinki (1964).

### Materials and procedure

#### Pre-scan session

Immediately before scanning, participants performed a series of daily-life activities at different locations on the campus of the University of Liège while wearing an Autographer (OMG Life Ltd., Kidlington, UK) tied around their neck. The Autographer is a small wearable camera that automatically and silently takes a continuous series of pictures from the first-person perspective through a 136° fish-eye lens, depending on several electronic sensors (e.g., temperature, accelerometer, magnetometer, etc.). This device has been used in recent neuroimaging and behavioral studies investigating episodic memories for real-life events.^36^ To record the entire set of events as completely as possible, the Autographer was set to the fastest capture rate (approximatively 10 pictures per minute).

The entire set of events was designed to simulate activities that college students would perform in their daily life. It consisted of two kinds of events: goal-directed actions at specific locations, and spatial displacements that did not involve any particular action other than walking from a place to another. These two kinds of events correspond respectively to what Tversky et al.^39^ referred to as “events by hands” (i.e., interactions with objects; e.g., buying a snack) and “events by feet” (i.e., changes of location; e.g., going from the laboratory building to the bus stop). We initially planned to examine a third kind of events that consisted in waiting at specific locations without doing any particular action (e.g., waiting at the bus stop); this condition was included as we assumed it would involve fewer perceptual changes than the two previous types of events. However, preliminary behavioral analyses (see Supplemental Information) revealed that this was not the case and that the variance in recalled elements (i.e., information about persons, objects, thoughts, and perceptual details) was much higher in memories for waiting moments than in memories for actions and spatial displacements, showing that the former condition did not involve a homogenous set of events (the waiting moments took place in public spaces, such as a bus stop, where many different events could occur while participants were waiting). Thus, we decided not to report analyses about waiting events in the main article because this heterogeneity in recalled contents make the results difficult to interpret, but these are available in the Supplemental Information.

Participants were instructed to perform the following series of successive actions: First, they left the laboratory, went to the bus stop, and stayed there as if they were waiting for the bus. After having waited approximately two minutes, they went to the campus shop to buy a snack of their choice. Next, they went to the elementary school near the University campus and stayed in front of the school for approximately two minutes as if they were waiting for a kid. After that, participants went to the Speech and Language Therapy Service to post a letter in a particular mailbox. Having posted the letter, they came back to the laboratory building to print a document and to wait in the laboratory building hall (again for approximately two minutes) as if they were waiting for an appointment with a professor. Then, they went to a cafeteria to buy a drink of their choice, after which they chose a table inside the cafeteria to wait as if they were waiting for a friend. Next, participants went to a classroom and waited outside as if they were waiting for the beginning of a course. Finally, they were instructed to return to the laboratory and answered an email sent by the experimenter during the walk.

Before starting the walk, participants were given the letter that they should post, as well as 5 euros for purchasing the snack and the drink. They were told that they could take the route they wanted to go from each location to the next, but not to perform any additional actions during spatial displacements or waiting moments (e.g., using their smartphones, listening to music, or counting down time). In addition, participants were asked to avoid obstructing the lens of the camera and to behave as naturally as possible. Finally, they were informed that pictures taken during their walk would be presented during a subsequent fMRI session, but no mention was made of the memory task. Instead, we used a cover story in which participants were told that they would have to perform aesthetic judgments on the pictures in the scanner. We used an incidental encoding paradigm because in everyday life memory encoding is most of the time an incidental process that happens without intentional control.^1^

#### fMRI Session

Immediately after the walk on campus, participants took part in an fMRI session that involved the recall of events experienced during this walk, as well as a control condition that was intermixed with the episodic memory conditions. The entire task comprised 80 trials (20 trials in each condition). The order of trial presentation was randomized with the constraint that two subsequent trials of the same condition were not presented in direct succession or with an interval of more than five trials.

##### Episodic memory tasks

In the episodic memory conditions (i.e., action, spatial displacement, and waiting conditions), participants were asked to remember the unfolding of actions, spatial displacements, or waiting periods. Each trial started with a fixation cross (with a variable duration ranging between one and three seconds) followed by the presentation of a written experimental instruction (for one second), specifying the task that participants should perform on the trial (i.e., remembering the unfolding of an event or remembering a particular moment of waiting). Then, a picture taken during the pre-scan walk of the participant was presented, which corresponded to the beginning of either a particular goal-directed action, a spatial displacement, or a waiting moment. Experimental instructions specified that each picture would be displayed several times during the fMRI session. In total, 20 trials were presented in each of the episodic memory conditions (five pictures that were each repeated four times).

For each picture, participants were first asked to identify the corresponding moment of the walk (without starting to remember the corresponding event); all pictures were presented to the participants before the scan session to ensure that they would be able to easily identify the corresponding moment of their walk during the fMRI session. As soon as they identified the moment depicted, participants were instructed to close their eyes and then to press a response key to indicate that they began their mental replay (their button press was confirmed by a sound signal). Depending on the condition, they were asked to mentally replay either the unfolding of the event (for actions and spatial displacements) or the waiting period, in as much detail as possible, starting at the precise moment they identified on the picture. Participants were told that their mental replay could be of anything that they encountered during the walk (e.g., people, objects, actions, thoughts, etc.). Moreover, in the unfolding conditions, it was specified that the task was not to proceed as far as possible in the recall of the walk or action, but rather to be as precise and detailed as possible. In the waiting condition, participants were told that they should only remember what they experienced during the moments in which they waited at a particular location (without remembering the unfolding of what happened after these waiting moments). The mental replay phase lasted 12 seconds after which a sound signal prompted participants to stop their mental replay and to open their eyes.

Following the mental replay phase, a one-second fixation cross was shown, immediately followed by the presentation of a four-point Likert scale assessing the extent to which participants’ mental replay was static (meaning that it was focused on a particular moment of past experience) or dynamic (meaning that it involved the unfolding of several moments of past experience; see Figure 1B). Responses were self-paced with a maximum of five seconds to respond.

##### Control task

In the control task (scene condition), participants had to form mental representations of static scenes. The structure and timing of the control condition was identical to the episodic memory conditions: Each trial started with a fixation cross, followed by written instructions. Then a picture of a scene was presented in the middle of the screen. Each picture represented a specific location on the campus of the University of Liège that participants did not visit during their walk (but that was taken with the same wearable camera). In total, five different pictures were presented four times each, resulting in 20 trials. As in the episodic memory conditions, each picture was also presented to the participants before the scan. After having examined the picture, participants were asked to close their eyes and then to press the response key to indicate that they began forming a mental representation of the scene (a sound signal confirmed their button press). They were instructed to maintain their mental representation of the scene in as much detail as possible until the presentation of a second sound signal (i.e., after 12 seconds). It was specified that their task was not to retrieve personal memories (e.g., past events experienced at the presented location or past experiences that happened at similar locations) but only to mentally represent the scene that was depicted on the picture. The control condition trials ended with the presentation of a four-point Likert scale assessing to what extent each trial involved a static representation or the unfolding of several moments (as in the episodic memory conditions). We decided to use pictures of scenes that were not encountered by participants during their tour for the control condition, as scenes that were experienced during the tour might be difficult to represent statically (i.e., the pictures might automatically trigger dynamic event recall).

#### fMRI data acquisition

All MRI data were acquired on a whole-body 3T scanner (Magnetom Prisma, Siemens Medical Solutions, Erlangen, Germany) operated with a 20-channel receiver head coil. Multislice T2*-weighted functional images were acquired using a gradient-echo-planar imaging (EPI) sequence (TR = 1170 ms, TE = 30 ms, FA 90°, matrix size 72 × 72 × 36, voxel size 3 × 3 × 3 mm^3^). Thirty-six 3 mm thick transverse slices (FOV 216 × 216 mm^2^) were acquired, with a 25% interslice gap, covering the whole brain. Around 1400 functional volumes were obtained for the entire task and we dropped the five initial volumes to avoid T1 saturation effect. After image acquisitions, we generated field maps using a double echo gradient-recalled sequence (TR = 634 ms, TE = 10 and 12.46 ms, FoV = 192 × 192 mm2, 64 × 64 matrix, 40 transverse slices with 3 mm thickness and 25% gap, flip angle = 90°, bandwidth = 260 Hz/pixel) that we used to correct echo-planar images for geometric distortion due to field inhomogeneities. Finally, we acquired a structural MR scan [T1-weighted 3D magnetization-prepared rapid acquisition gradient echo (MP-RAGE) sequence, TR = 1900 ms, TE = 2.19 ms, FOV 256 × 240 mm2, matrix size 256 × 240 × 224, voxel size 1 × 1 × 1 mm]. All experimental stimuli were presented on a screen positioned at the rear of the scanner, which the subject could see through a mirror mounted on the head coil.

#### Post-scan debriefing

Immediately after the fMRI session, participants were shown the 20 pictures presented during the scanning session (i.e., 5 pictures per condition) and, for each picture, they were asked to describe the content of the mental representation they formed in the scanner (i.e., their memories or scene representations). Verbal reports were recorded using a digital audio recorder. Furthermore, in the episodic memory conditions, each verbal report was followed by a review of the pictures that had been taken during the walk and participants were instructed to select the picture that best corresponded to their mental representation of the last moment they remembered while mentally replaying the unfolding of the event in the scanner (i.e., the moment they remembered when hearing the second sound signal). These tasks allowed us to determine (a) whether the participants correctly performed all conditions in the scanner and (b) to estimate the density of experience units recalled (see Behavioral data analyses). Note that while this measure allowed us to confirm that the memory conditions differed in terms of the density of recalled experience units, it only provides a rough estimate of the density of recalled units at the trial level because we only have one measure of unit density per event (i.e., each event was repeated four times in the scanner, but we only obtained one density measure per event in the post-scan session). For this reason, this measure was not used as a parametric regressor in the fMRI analyses.

Finally, participants were asked to rate the overall clarity of the memory or scene they formed in response to each picture (from 1 = *not clear at all*, to 7 = *extremely clear*), as well as the easiness of remembering/imagining (from 1 = *not easy at all*, to 7 = *extremely easy*). Furthermore, for trials of the episodic memory conditions, participants were asked to estimate their overall feeling of re-experiencing the unfolding of events (from 1 = *not at all*, to 7 = *completely*).

#### Behavioral data analyses

Verbal descriptions of the memories reported during the post-scan interview consisted of a succession of distinct moments of past experience—referred to as experience units.^12,13,17^ For example, a typical verbal report would start by “I got out of the building” (first experience unit), “then I walked along the parking” (second experience unit), “then, I saw a man I know on the opposite side of the road” (third experience unit), and so on. The segmentation of verbal reports into distinct experience units was performed by the first author based on indications of transitions or temporal discontinuities between reported moments of experience (i.e., verbal clues such as “then”, “next”, and “after that” or moments of silence); most of the time, transitions between experience units involved significant changes in actions, environmental elements, and/or thoughts (e.g., “I saw a sheet of paper on the ground just in front of the door of the cafeteria,” followed by “Once inside of the cafeteria, I was happy to see that there were few students queuing”). Although in most cases this segmentation procedure was obvious, we nevertheless assessed its reliability by asking a second trained rater to independently segment a random selection of 20% of verbal reports. The Intraclass Correlation Coefficient (ICC) computed on the number of experience units that were identified within verbal reports showed an excellent agreement between the two raters (ICC = 0.91).

In addition to computing the number of experience units reported by participants, we estimated the density of recalled experience units in the two episodic memory conditions by using temporal information associated with the pictures acquired during the walk. More specifically, for each trial, we estimated the actual event duration by calculating the amount of time between the beginning of each event and the end of the event. The beginning corresponded to the picture presented on the computer screen during each fMRI trial. The end of the event corresponded to the picture selected by participants as the last moment of the experience they remembered in the scanner. Then, we estimated the density of recalled experience units as the number of experience units reported per minute of the actual event duration.^12,17^

Behavioral data were analyzed using robust statistical methods (i.e., Yuen’s *t* test)^65^ because the assumptions underlying classical inferential methods (normality and homoscedasticity) were violated for our main measures of interest (i.e., the number of reported experience units per minute of the actual event duration). These methods perform well in terms of type I error control and statistical power, even when the normality and homoscedasticity assumptions are violated.^66^ Effect sizes were estimated using the explanatory measure of effect size *ξ*: values of 0.10, 0.30, and 0.50 correspond to small, medium, and large effect sizes, respectively. All descriptive statistics refer to the 20% trimmed means and their 95% confidence intervals. These analyses were conducted in R^67^ using the functions provided by Wilcox.^68^

#### fMRI data preprocessing

Image preprocessing was performed using the SPM 12 software (Wellcome Department of Imaging Neuroscience, http://www.fil.ion.ucl.ac.uk/spm) implemented in MATLAB R2017a. EPI time series were corrected for motion and distortion artefacts using Realign and Unwarp^69^ with the Fieldmap Toolbox.^70^ The mean realigned EPI image was coregistered with the structural T1 image, and the coregistration parameters were applied to the realigned EPI time series. The T1 image was segmented into gray matter, white matter, and cerebrospinal fluid, using the unified segmentation approach,^71^ and we used the normalization parameters obtained from the segmentation procedure to normalize the functional images to MNI space (2×2×2 mm voxels). Finally, we performed a spatial smoothing of functional images using an 8 mm FWHM Gaussian kernel.

For the representational similarity analyses, slice time corrected volumes coregistered to the first scan of each subject were used to perform the searchlight in the native space of each subject. To exclude variance explained by motion, regressors from motion correction were used as predictors in a GLM for the voxel time series in every voxel. The resulting residuals were then used as the final input for the analyses. To exclude that the searchlight results are influenced by condition differences in motion inside the MRI we analyzed the motion parameters resulting from fMRI preprocessing. As the assumptions for an analysis of variance (ANOVA) were violated we evaluated motion effects with a non-parametric permutation test using ez-Perm^72^ in R.^67^ No significant differences of the motion regressor between the three conditions were found.

#### fMRI data analyses

##### Univariate analyses

Univariate analyses were performed using SPM 12. For each participant, BOLD responses were modeled at each voxel, using a general linear model (GLM). The memory/scene representation phase of each trial was modeled separately for each condition (i.e., action, spatial displacement, waiting, and scene representation) as epoch-related responses (beginning at the key press and ending at the second sound signal after 12 seconds) and convolved with the canonical hemodynamic response function to create the regressors of interest. Trials for which participants did not press the response button to indicate that they identified the picture and trials for which ratings indicated that they did not perform the requested task were modelled in a separate regressor of no interest. This corresponds to trials for which they provided a rating of 1 or 2 in the two episodic memory conditions (indicating that they mentally represented only one single moment of past experience), or a rating of 3 or 4 in the scene condition (indicating that they mentally represented the unfolding of several moments of past experience). Note that more than 80% of trials in the episodic memory conditions received a rating of 4, and more than 90% of trials in the control condition a rating of 1, so that there was not enough variance to include the ratings as a parametric regressor in the analyses. Experimental phases corresponding to the presentation of the pictures and the presentation of the Likert scales were also modelled as epoch-related responses with two regressors of no interest (i.e., a single regressor for each phase across all conditions). Finally, we also modelled the two beeps presented respectively at the beginning and the end of each task as an event-related response, with a single regressor across all conditions. Finally, the design matrix included the realignment parameters to account for any residual movement-related effect. The canonical hemodynamic response function was used and a high-pass filter was implemented using a cutoff period of 128 seconds to remove low-frequency drifts from the time series.

A series of linear contrasts was computed to identify the brain regions involved in the processing of episodic versus scene representations, and in the retrieval of actions versus spatial displacements. More specifically, we first investigated brain regions showing higher activity during each episodic memory condition compared to scene representation (i.e., Action > Scene; Spatial displacement > Scene). Next, we examined brain regions showing differential activity during the retrieval of the goal-directed actions and spatial displacements (Action > Spatial displacement, and vice versa).

The contrasts of interest were first computed for each participant and were then entered into random effects one-sample *t*-tests. For all analyses, statistical parametric maps (SPMs) were thresholded at *p* < .001, uncorrected, and statistical inferences were corrected for multiple comparisons at the voxel level (*p* < .05, FWE-corrected) using a small volume correction (SVC) within a priori regions of interest. These a priori regions of interest corresponded to brain areas that have been previously related to autobiographical memory, as determined by a meta-analysis of 84 studies performed with Neurosynth (www.neurosynth.org).^73^ These regions were merged into a single mask for the SVC (8156 voxels). The coordinates reported in Tables 2 and 3 refer to brain regions identified in the SPMs thresholded at *p* < .001 (uncorrected) that were statistically significant when the SVC was applied. Only clusters with a size of k > 15 voxels are reported, as small clusters of above threshold voxels can be the result of spatial smearing (due to data smoothness introduced during preprocessing) around isolated voxels with high *t*-values. For visualization purposes, Figure 2 displays activations thresholded at *p* < .001 (uncorrected) that fell within the autobiographical network mask.

##### Representational Similarity Analysis

In addition to the univariate analyses, we conducted a representational similarity analysis, a well-established method to study neural representational structures in brain activity patterns.^74^ First, we used this method to assess whether the different conditions (i.e., action, spatial displacement, and scene representation) showed distinct patterns of voxel activity during the recall period. This would substantiate further analysis of the temporal dynamics of multi-voxel pattern activity (see below). We computed a single-trial GLM for each subject with one regressor for each recall period of 12 seconds, resulting in 60 beta weights for each subject. Within the autobiographical network mask (as used for the univariate analyses), we tested whether trials within one condition were more similar to each other compared to trials of other conditions. We calculated the cross-correlation between all trials using spearman rank correlations, resulting in a 60 × 60 matrix. Correlation coefficients between all pairs of trials in each condition were used to build the mean within-condition correlation coefficient. Correlations of each trial with itself were excluded before computing the mean. To generate between-condition correlation coefficients, trials of one condition were correlated to trials of the other conditions. In addition, we correlated each recall trial beta to its respective image cue beta, which we derived from a single-trial GLM for each subject with one regressor for each cue period. The resulting 60 recall x 60 cue image correlation matrix was then used to calculate the correlation coefficients of each recall event to its own cue and the cue of other trials of the same event (own event) and to cues of other events from the same condition (other events). Differences for within and between condition/event similarities were analyzed using a using a repeated measures ANOVA implemented in the ez-package^72^ in R.^67^ For significant effects, we report Cohen’s *d* as measure of effect size.^75^

##### Analysis of temporal dynamics

Next, we investigated the temporal dynamics of patterns of brain activity during the memory/scene construction period by computed correlations between consecutive MRI volumes during the recall phase. We expected dynamic and high-density events to show higher differences in brain activity patterns across time, as indicated by smaller volume to volume correlations. The analysis was limited to grey matter voxels by applying a subject specific grey matter mask segmented using Freesurfer (version 6).^76^ We constructed a sphere with a radius of 6 voxels which was centered at each voxel. If the sphere consisted of fewer than 20 voxels no further analysis was performed at this location. At each valid sphere location, 10 consecutive volumes from the recall period, starting with the fifth volume after the recall onset to account for the delay of the hrf, were used to compute a 10×10 crosscorrelation matrix (Figure 4A). To analyze consecutive changes in voxel patterns, the first off-diagonal was used to extract Spearman’s rank correlations of each volume with the next (i.e., volume 5 was correlated with volume 6; volume 6 with volume 7; etc.). Then, we computed the mean over the first off-diagonal for the resulting 9 volume-to-volume correlations. This was done for each trial in each condition. In a final step, the mean over all fisher-z-transformed volume-to-volume correlations in all trials per condition was saved at the current sphere location. Thus, three correlation coefficient maps, one per condition, were created for each subject. For group level analysis, the subject-wise correlation coefficient maps were normalized to the FSL standard 2mm 152 MNI template using a non-linear registration in FSL (version 5.0.9, http://fsl.fmrib.ox.ac.uk/fsl/fslwiki/).

On the group level, the correlation coefficient maps were used to identify regions with consistent differences in correlations between the conditions. Using *randomise* in FSL,^77^ a non-parametric version of the paired one-sample *t*-test for each condition pairing was conducted. *Randomise* analysis was performed using 10,000 random sign-flips and threshold-free cluster enhancement (TFCE)^78^ was used to identify significant clusters. TFCE has been shown to be more sensitive than clusterwise thresholding methods by taking both voxelwise signal strength and extent of neighboring voxels into account without relying on an absolute threshold.^78^ All results reported are FWE corrected based on TFCE and thresholded at *p* < 0.05 to control for multiple comparisons. We used the Harvard-Oxford Cortical Structural atlas^79^ for anatomical labels.

## SUPPLEMENTAL INFORMATION

### Supplemental behavioral results

In addition to computing the number of recalled experience units based on verbal descriptions of memories obtained after the scanning session (see STAR Methods), we also scored the content of these memories using a scoring system developed in previous behavioral studies.^1^ Each recalled element was classified using the following categories: person, object, thought, action with interaction, displacement, spatial detail, and perceptual detail (see **Table S1** for descriptions and examples of each category).

**Table S1.**
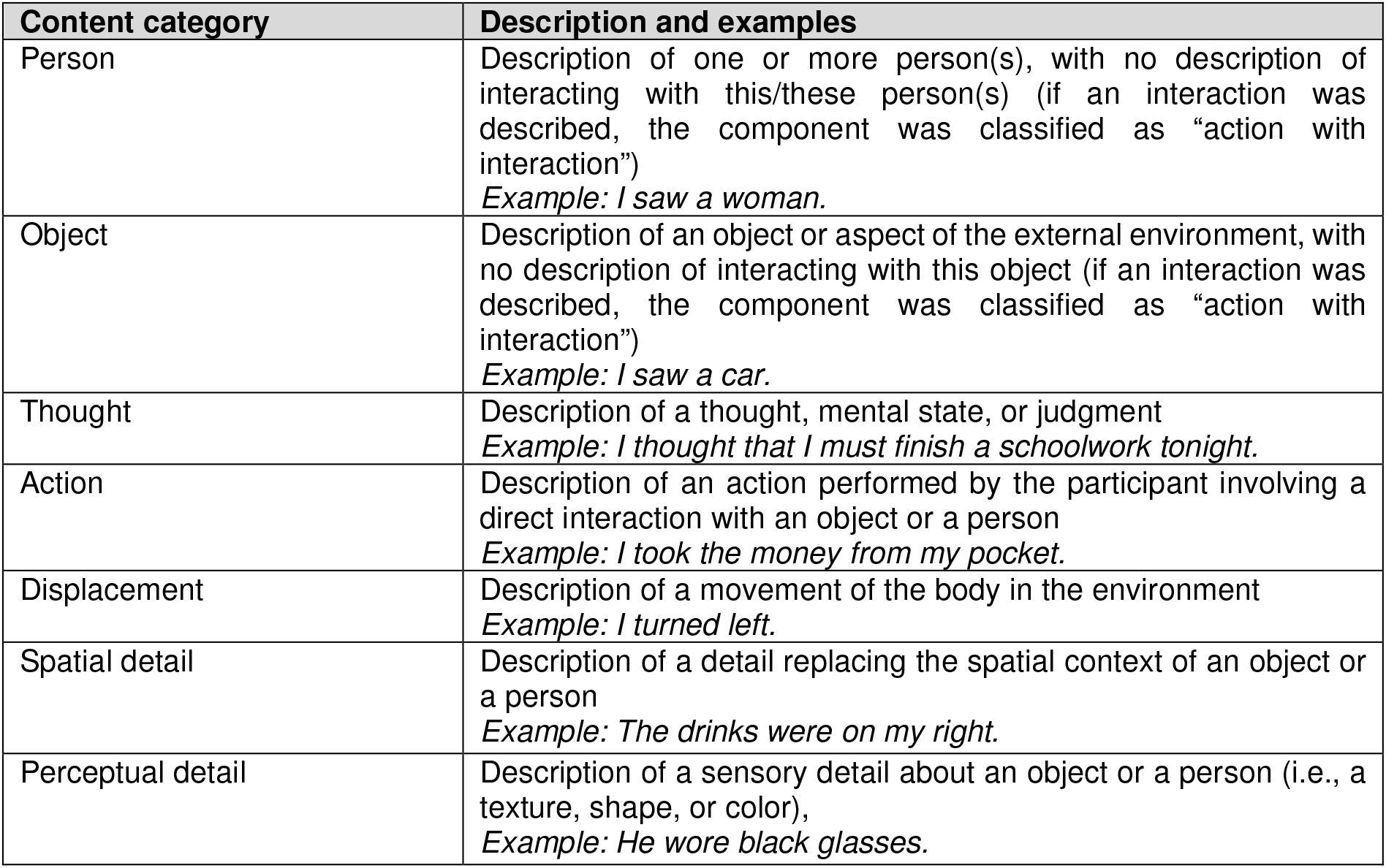
Descriptions and examples of content categories

| Content category | Description and examples |
| --- | --- |
| Person | Description of one or more person(s), with no description of interacting with this/these person(s) (if an interaction was described, the component was classified as “action with interaction”)<br>*Example: I saw a woman.* |
| Object | Description of an object or aspect of the external environment, with no description of interacting with this object (if an interaction was described, the component was classified as “action with interaction”)<br>*Example: I saw a car.* |
| Thought | Description of a thought, mental state, or judgment<br>*Example: I thought that I must finish a schoolwork tonight.* |
| Action | Description of an action performed by the participant involving a direct interaction with an object or a person<br>*Example: I took the money from my pocket.* |
| Displacement | Description of a movement of the body in the environment<br>*Example: I turned left.* |
| Spatial detail | Description of a detail replacing the spatial context of an object or a person<br>*Example: The drinks were on my right.* |
| Perceptual detail | Description of a sensory detail about an object or a person (i.e., a texture, shape, or color),<br>*Example: He wore black glasses.* |

The mean number of components included in the three types of memories (i.e., actions, spatial displacements, and waiting moments) are shown in **Table S2**. As expected, memories for actions contained more descriptions of actions, *t* = 10.91, *p* < 0.001, *ξ* = 0.93, whereas memories for spatial displacements contained more descriptions of displacements, *t* = 2.39, *p* = 0.001, *ξ* =0.92. There was no significant difference between these two types of memories for the other components, except for thoughts, *t* = 3.37, *p* = .006, *ξ* =0.42.

Contrary to our expectations, memories of waiting moments did not include less perceptual detail (in fact, they included numerically more detail; see **Table S2**) than the other two types of memories (*t* = 0.72, *p* = 0.487, *ξ* =0.18 and *t* = 2.13, *p* = 0.054, *ξ* =0.61 for comparisons with actions and spatial displacements, respectively). Furthermore, memories of waiting moments included significantly more descriptions of persons than the other two types of memories (*t* = 3.44, *p* = 0.005, *ξ* =0.67 and *t* = 4.26, *p* = 0.001, *ξ* =0.69 for comparisons with actions and spatial displacements, respectively). In addition, we examined the variance in recalled contents across memories within each condition and found that the variance in several recalled elements (i.e., persons, objects, thoughts, and perceptual details) was higher in memories of waiting moments than in memories of actions and spatial displacements (see **Table S3**), showing that the former condition involved a more heterogeneous set of events.

**Table S2.**
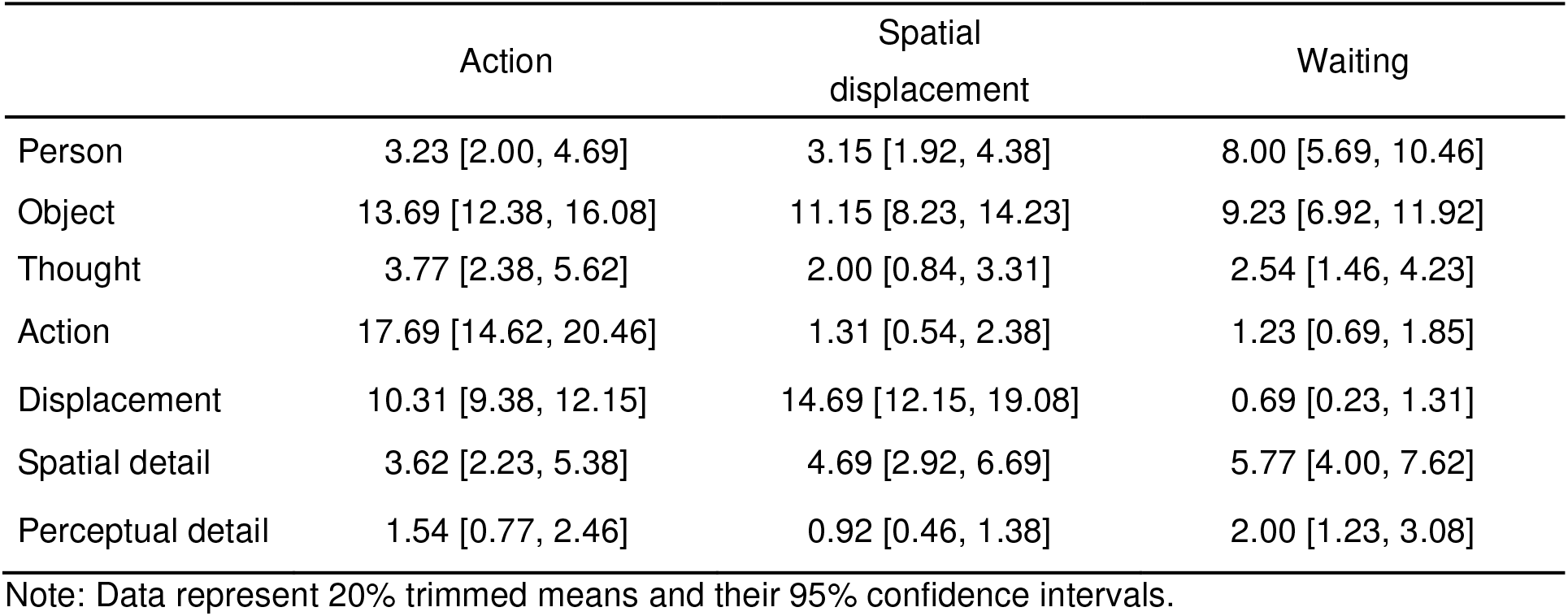
Components recalled in the three memory conditions

**Table S3.** Variance in recalled contents across memories in the three conditions

|  | Action | Spatial displacement | Waiting |
| --- | --- | --- | --- |
| Person | 0.018 | 0.034 | 0.266 |
| Object | 0.045 | 0.090 | 0.168 |
| Thought | 0.09 | 0.018 | 0.045 |
| Action | 0.055 | 0.018 | 0.019 |
| Displacement | 0.044 | 0.063 | 0.008 |
| Spatial detail | 0.019 | 0.052 | 0.077 |
| Perceptual detail | 0.005 | 0.007 | 0.032 |

Note: For each memory, the proportion of each recalled content per unit of experience was computed. The cells show the variance of these within each memory condition.

### Supplemental fMRI results

#### Univariate activity associated with waiting moments vs scene representations

We conducted a univariate analysis (see STAR Methods) to identify brain regions within the autobiographical memory network that showed higher activity when recalling waiting moments compared to scene representations. Results showed that the recall of waiting moments was associated with increased activation in medial parietal areas (including the precuneus, posterior cingulate, and retrosplenial cortex), angular gyrus, middle frontal gyrus, middle temporal gyrus, middle occipital cortex, supplementary motor area, and cerebellum (see **Table S4**).

**Table S4.** Brain regions showing increased activity for waiting memories vs scene representations

| Brain regions | Side | MNI coordinates |  |  | Cluster Size<br>(n° voxels) | *t* |
| --- | --- | --- | --- | --- | --- | --- |
|  |  | X | Y | Z |  |  |
| Precuneus/PCC/RSC | L/R | 6 | -68 | 38 | 2104 | 12.82 |
|  |  | 10 | -66 | 32 |  | 10.47 |
|  |  | -4 | -66 | 26 |  | 11.04 |
|  |  | -10 | -60 | 20 |  | 9.52 |
| Middle frontal gyrus | L | -38 | 10 | 56 | 91 | 8.14 |
|  | L | -24 | 18 | 48 | 71 | 4.62 |
|  | R | 28 | 20 | 54 | 28 | 5.60 |
| AG | R | 42 | -56 | 24 | 215 | 6.91 |
| Middle occipital gyrus | L | -40 | -72 | 38 | 52 | 6.25 |
|  | L | -44 | -72 | 36 | 26 | 5.08 |
| Cerebellum | R | 22 | -76 | -28 | 59 | 5.31 |
| MTG | L | -52 | -56 | 18 | 29 | 4.85 |
|  | L | -58 | -56 | 22 | 16 | 4.80 |
| SMA | L | -8 | 16 | 62 | 27 | 4.75 |

**Note:** Coordinates refer to brain regions identified in the SPMs thresholded at $p < .001$ (uncorrected) that were statistically significant (at $p < .05$ ) using SVC (see STAR Methods). PCC = Posterior cingulate cortex; RSC = Retrosplenial cortex; MTG = Middle temporal gyrus; AG = Angular gyrus; SMA = supplementary motor area.

#### Patterns of brain activity in the waiting condition

We initially conducted a representational similarity analysis (see STAR Methods) to analyze content-specific representations between and within trials of the four conditions. When including the waiting condition, a repeated measures ANOVA revealed higher similarities within the same condition compared to between condition trials (*F*(1,20) = 36.65, *p* = <0.0001). There was no effect of condition (action/spatial displacement/control/waiting; *F*(3,60) = 1.09, *p* = 0.359) and no interaction (*F*(3,60) = 2.00, *p* = 0.123; **Figure S1A**).

Furthermore, we analyzed whether neural activity patterns of the same events were more similar to their own image cues compared to image cues of other events in the same condition (**Figure S1B**). Since similarities within trials tend to be higher than between trials, we calculated, for each event, neural activity during the recall phase in trial m with activity during the image cues in trials n≠m of the same event and compared this to correlations with the image cues of the other events in the same condition. When including the waiting condition, a repeated measures ANOVA showed a significant main effect of cue type (own vs. other event; *F*(1,20) = 48.84, *p* = <0.0001), no main effect of condition (*F*(3,60) = 0.99, *p* = 0.388), and no interaction (*F*(3,60) = 2.29, *p* = 0.09).

**Figure S1.**
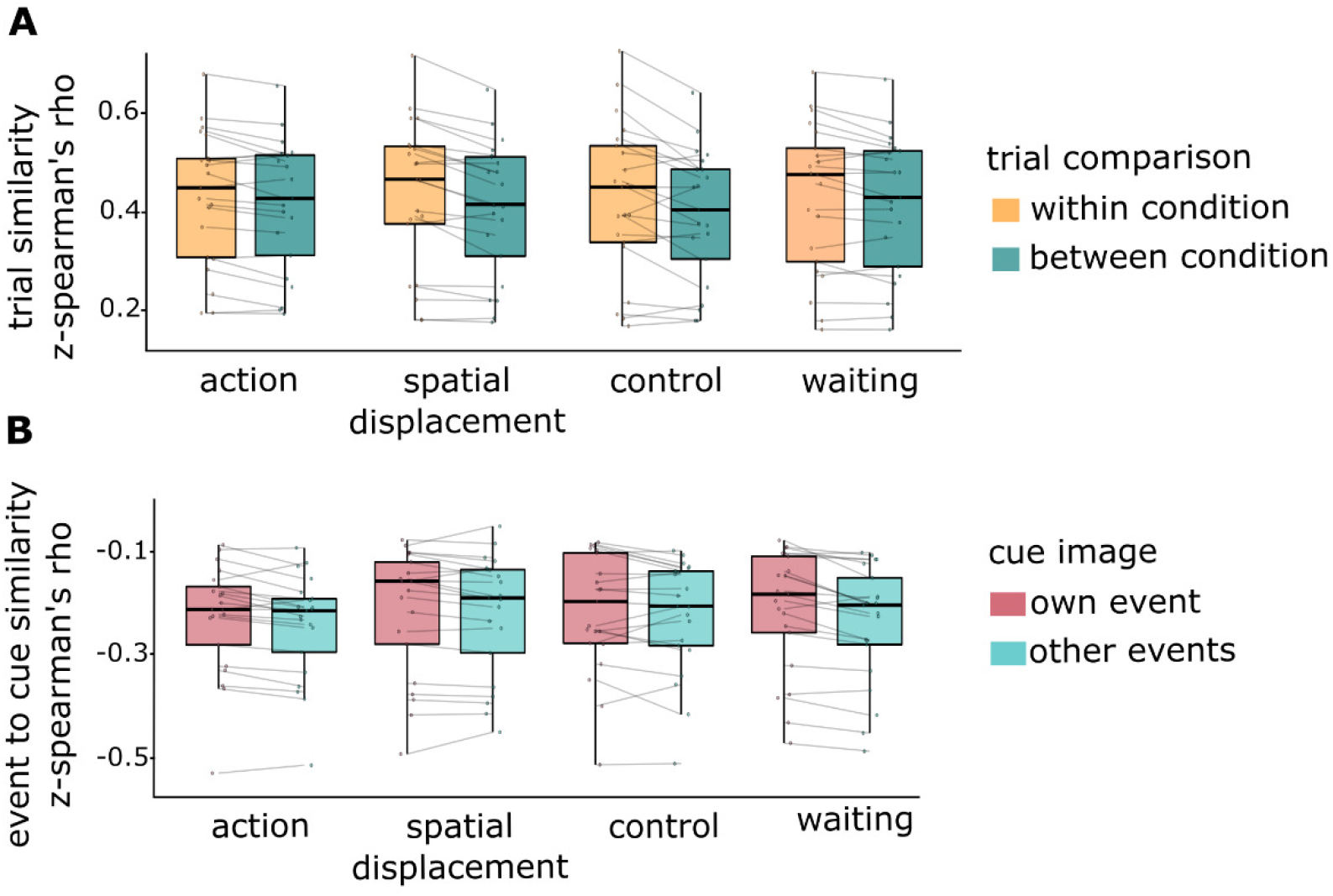
Content-specific memory representations in the autobiographical memory network. (A) Distributed patterns of neural activity are more similar when recalling events from the same condition than events from different conditions. This effect is observed across all experimental conditions. (B) Neural patterns of action and spatial displacement recall are more similar to their trial-specific image cue than to image cues of other events of the same condition. This effect is found across conditions. Dots show descriptive means for each subject; individual observations are linked with lines.

#### Patterns of brain activity across repetitions

All events were recalled four times to achieve a sufficient number of trials for multivariate pattern analyses. To investigate whether differences between conditions might have been caused solely by these repetitions, we conducted an additional representational similarity analysis (using the same autobiographical network mask as for the univariate analyses). If repeating trials had an effect on differences of neural activity patterns between conditions, we would expect an interaction between condition and repetitions. To investigate this possibility, we compared the similarities of each trial to all other repetitions of the same event. Recall period beta weights derived from a single-trial GLM were used to correlate each event trial to the other three repetitions of the same event. We thereby obtained one similarity measure for each trial by averaging the correlation coefficients of this trial to the other three trials of the same event (i.e., how similar is the neural activation pattern of trial 2 event A to the pattern of trial 1, 3 and 4 of event A). This similarity value thus reflected the overall similarity of a trial to all its repetitions. Using a repeated-measures ANOVA, we found a main effect of repetition (*F*(3, 60) = 5.32, *p* = 0.0044, ηp^2^ = 0.05), but no effect of condition (*F*(3, 60) = 1.54, *p* = 0.23) and no interaction (*F*(9, 180) = 0.47, *p* = 0.77). Post-hoc t-tests further investigating the main effect of repetition revealed that the last repetition (*M* = 0.41, *SD* = 0.16) showed a significantly lower similarity to all other repetitions (1^st^ : *t*(83) = 3.85, *p_bonferroni_* = 0.0010, *d* = 0.42; 2^nd^: *t*(83) = 3.84, *p_bonferroni_* = 0.0010, *d* = 0.41; 3^rd^ : *t*(83) = 4.13, *p_bonferroni_* = 0.0003, *d* = 0.47) (**Figure S2**), whereas first (*M* = 0.43, *SD* = 0.15), second (*M* = 0.44, *SD* = 0.16) and third (*M* = 0.44, *SD* = 0.16) repetitions did not significantly differ. This suggests that neural activity patterns of an event repeatedly recalled only began to decrease in similarity during the last repetition. However, this effect of repetitions did not differ between conditions (see **Figure S2**). Importantly, since we did not find an interaction, repeating recall phases could not explain the condition differences between conditions.

**Figure S2.**
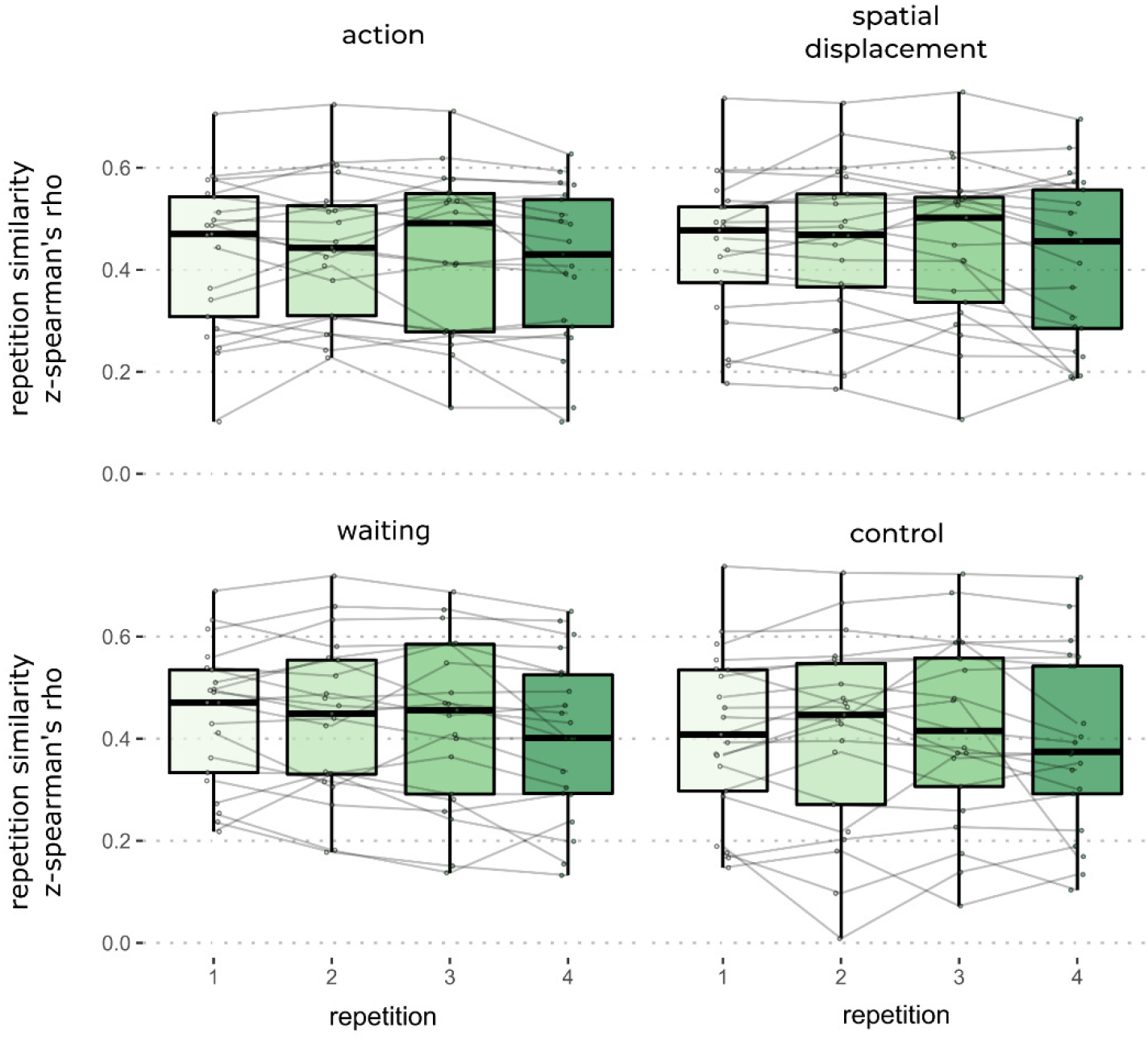
Patterns of brain activity across repetitions. Distributed patterns of neural activity across repetitions did not differ between conditions. Dots show descriptive means for each subject; individual observations are linked with lines.

#### Temporal dynamics of brain activity patterns in the waiting condition

The behavioral results described above showed that the waiting condition involved a less consistent set of events, hence our decision not to report them in the main text. Nevertheless, for the sake of completeness, we also performed a searchlight analysis comparing the temporal dynamics of patterns of brain activity during the waiting recall period and the recall period of the other conditions (**Figure S3**). This analysis revealed higher temporal dynamics (i.e., lower autocorrelations) during the recall of waiting events compared to the control condition (static scenes) in a cluster including the frontal pole, the middle frontal gyrus, the lateral occipital cortex, and the precuneus (*t*(20) = 8.76; *p*_corr_ < 0.05, FWE corrected using TFCE). In comparison to spatial displacements, the recall of waiting episodes was associated with higher temporal dynamics (i.e., lower autocorrelations) in a cluster including the lateral occipital cortex, the posterior cingulate gyrus, the precuneus, cuneus, and the lingual gyrus (*t*(20) = 8.15; *p*_corr_ < 0.05, FWE corrected using TFCE).

**Figure S3.**
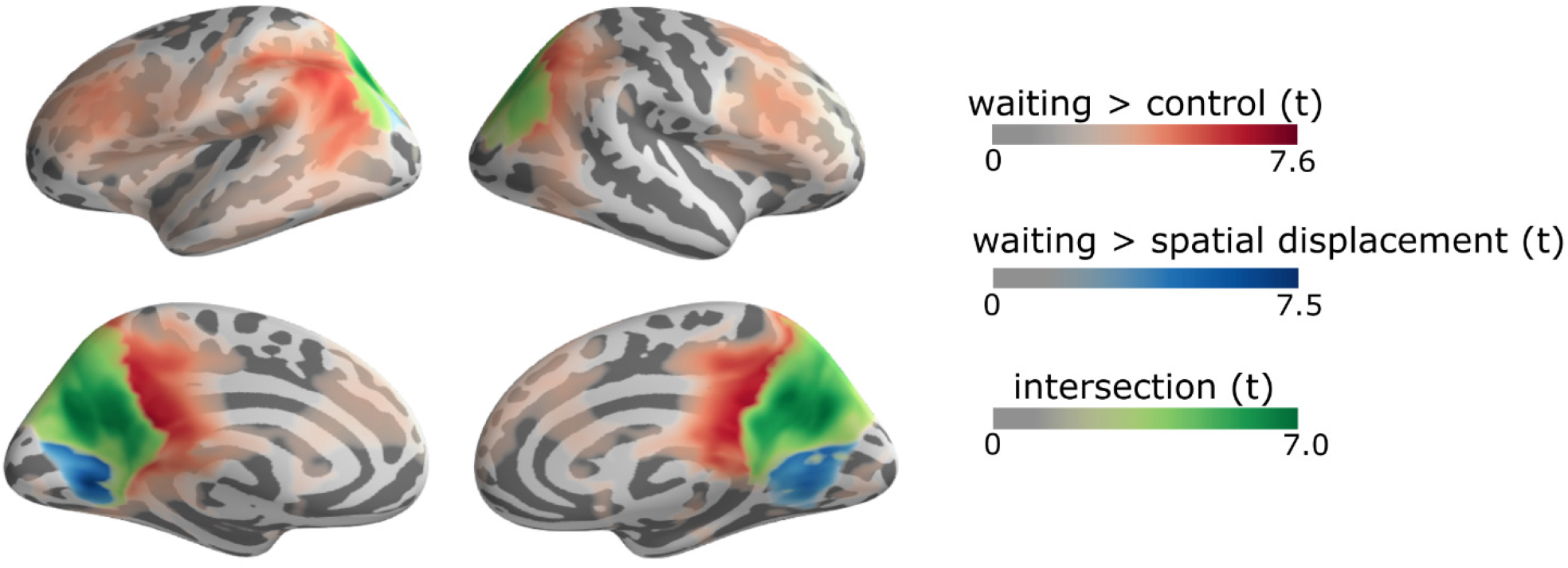
Temporal dynamics of patterns of brain activity during the recall of waiting periods. The searchlight analysis revealed higher temporal dynamics for the waiting condition compared to the control and spatial displacements conditions. The conjunction of higher temporal dynamics for waiting compared to both the spatial displacements and control conditions is shown in green. There was no significant difference between temporal dynamics of the waiting and action conditions.

## Footnotes

a Note that we refer to the memories of real-life events that we investigated in this study as episodic rather than autobiographical because the events recalled were very recent and, according to the influential view of autobiographical memory proposed by Conway,^19^ these memories only become truly autobiographical when they are integrated with autobiographical knowledge structures (i.e., general knowledge about the self and one’s personal life), so that they are retained in a durable form. However, this terminological distinction between episodic and autobiographical memories is not central to the main purpose of our study, which was to investigate the neural mechanisms underlying the dynamic unfolding of events within memory representations.

